# DDX3 is exploited by Arenaviruses to suppress type I interferons and favor their replication

**DOI:** 10.1101/224725

**Authors:** María Eugenia Loureiro, Andre Luiz Zorzetto-Fernandes, Sheli Radoshitzky, Xiaoli Chi, Simone Dallari, Nuha Marooki, Psylvia Lèger, Sabrina Foscaldi, Sonia Sharma, Nora López, Juan Carlos de la Torre, Sina Bavari, Elina Zúñiga

**Affiliations:** Division of Biological Sciences, University of California San Diego, La Jolla, CA, USA.; Molecular and Translational Sciences Division, United States Army Medical Research Institute of Infectious Diseases, Frederick, MD, USA.; La Jolla Institute for Allergy and Immunology, La Jolla, CA, USA.; Centro de Virología Animal, Instituto de Ciencia y Tecnología Dr. César Milstein, Consejo Nacional de Investigaciones Científicas y Técnicas, Buenos Aires, Argentina.; The Scripps Research Institute, Department of Immunology and Microbiology, La Jolla, CA, USA.

**Author notes:** Current address: Centro de Virología Animal, Instituto de Ciencia y Tecnología Dr. César Milstein, Consejo Nacional de Investigaciones Científicas y Técnicas, Buenos Aires, Argentina.

## Abstract

Several arenaviruses cause hemorrhagic fever (HF) diseases that are associated with high morbidity and mortality in humans. Accordingly, HF arenaviruses have been listed as top-priority emerging diseases for which countermeasures are urgently needed. Because arenavirus nucleoprotein (NP) plays critical roles in both virus multiplication and immune-evasion, we used an unbiased proteomic approach to identify NP-interacting proteins in human cells. DDX3, a DEAD-box ATP-dependent-RNA-helicase, interacted with NP in both NP-transfected and virus-infected cells. Importantly, DDX3 deficiency compromised the propagation of both Old and New World arenaviruses, including the HF arenaviruses Lassa and Junin viruses. The DDX3 role in promoting arenavirus multiplication correlated with both a previously un-recognized DDX3 contribution to type I interferon suppression in arenavirus infected cells and a positive effect of DDX3 on viral RNA synthesis. Our results uncover novel mechanisms used by arenavirus to exploit the host machinery and subvert immunity, singling out DDX3 as a potential host target for developing new therapies against highly pathogenic arenaviruses.

**AUTHOR SUMMARY:** Arenaviruses include severe clinical pathogens causing hemorrhagic fevers and have been recently incorporated by the World Health Organization in a list of critical emerging diseases for which additional research and identification of clinical targets is urgently required. A better understanding of how viral proteins interact with host cellular factors to favor arenavirus multiplication can illuminate novel pipelines on therapeutic strategies. Here we demonstrated that the ATP-dependent RNA helicase DDX3 interacted with the arenavirus nucleoprotein, which displays fundamental functions in different steps of the viral-cycle. Our work also revealed an unexpected new biology on the role that DDX3 might play during viral infections. In sharp contrast to previous studies showing DDX3 enhancement of IFN-I induction, we demonstrated that DDX3 suppressed IFN-I production at late time points after arenavirus infection, contributing to a DDX3 pro-viral effect. We also showed that early after infection, DDX3 pro-viral role was IFN-I independent and was mediated by DDX3 facilitation of viral RNA synthesis without affecting RNA translation. Altogether, our study established DDX3 as a critical host interacting partner of the arenavirus nucleoprotein and demonstrated two previously unrecognized DDX3-dependent strategies by which these deadly viruses exploit the host cellular machinery and suppress immunity.

## INTRODUCTION

Arenaviruses include highly pathogenic hemorrhagic fever (HF) viruses endemic to West Africa and South America. Lassa virus (LASV), is an Old World (OW) arenavirus highly prevalent in West Africa where it causes about 300,000 infections and > 5,000 deaths yearly due to Lassa fever (LF), with mortality rates rising up to 50% for hospitalized patients in some outbreaks and to 90% for women in the last month of pregnancy [1,2]. Notably, increased travelling has resulted in the importation of LF cases to Europe and United States, underscoring the global risk represented by this virus [3]. Likewise, several New World (NW) arenaviruses including Junin (JUNV), Machupo (MACV), Guanarito and Sabia, as well as the more recently reported Whitewater Arroyo and Chapare viruses, cause human hemorrhagic fevers with ~30% mortality [4]. In addition, mounting evidence indicates that lymphocytic choriomeningitis virus (LCMV), a globally distributed OW arenavirus, is a neglected human pathogen that causes congenital defects and poses a special threat to immunocompromised individuals [5,6]. The live-attenuated vaccine Candid#1 has been shown to be effective against Argentine HF caused by JUNV [7], but Candid#1 is only licensed in Argentina and it does not protect against other HF arenaviral diseases. There are no other licensed arenavirus vaccines and, with the exception of the treatment with immune plasma that is restricted to JUNV infections in endemic areas [8], anti-arenaviral therapy is limited to an off-label use of the nucleoside analog ribavirin that is only partially effective [9]. Accordingly, the World Health Organization (WHO) recently included Arenaviral HF in a list of emerging diseases for which additional research and identification of clinical targets are urgently needed [10]. A better understanding of how viral proteins interact with host cellular factors to enable arenavirus propagation could aid this task.

Arenaviruses are enveloped viruses with a negative-sense RNA genome, consisting of two single-stranded segments named S (ca. 3.4 kb) and L (ca. 7.2 kb). The nucleoprotein (NP), encoded by the S segment, is the most abundant viral protein and plays critical roles in different steps of the arenavirus life cycle [4,11–14]. In addition, the arenavirus NP counteracts the host type I interferon (IFN-I) response during viral infection by preventing the activation and nuclear translocation of interferon regulatory factor 3 (IRF-3), and subsequent induction of IFN-I production [15,16]. Arenavirus-NP has also been shown to inhibit nuclear translocation and transcriptional activity of NF-κB [17]. The anti-IFN-I activity of arenavirus NP was mapped to its C-terminal region and was associated to a folding domain corresponding to a functional 3’-5’ exonuclease of the DEDDH family [18,19]. NP has also been shown to interact with IKKε [20], MDA5 and RIG-I [21] which are involved in the IRF3 and NF-κB signaling pathways [22]. Moreover, it was recently demonstrated that NP can also subvert immune responses by associating to the dsRNA-activated protein kinase (PKR), a well-characterized antiviral protein that inhibits cap-dependent protein translation initiation via phosphorylation of eIF2α [23]. Thus, the arenavirus NP plays essential roles in viral multiplication and the virus’s ability to counteract key components of the host’s antiviral innate immune response. Targeting NP-host cell protein interactions required for NP to execute its functions could facilitate novel strategies to curtail arenavirus life cycle.

In the present study, we pursued an unbiased approach to identify novel host factors targeted by the arenavirus NP that could contribute to viral multiplication or be exploited by the virus to subvert the immune response. We found that DDX3, a DEAD (Asp-Glu-Ala-Asp)-box ATP-dependent RNA helicase, directly interacted with LCMV NP and was critical for supporting optimal LCMV growth, a finding that was extended to both OW and NW arenavirus infections in human cells. Strikingly, and in contrast to roles previously ascribed to DDX3 in promoting IFN-I [24,25], we observed that DDX3 contributed to IFN-I suppression upon arenavirus infection, partially explaining its pro-viral effect late in infection. In contrast, the early pro-viral effect of DDX3 was IFN-I independent and was explained by DDX3 positive effect on viral RNA synthesis. Our results uncovered previously unrecognized maneuvers evolved in highly pathogenic arenaviruses to favor their own growth by exploiting the host machinery and evading the immune system, raising DDX3 as a potential universal target for the rational design of antiviral therapies against arenaviruses infections.

## RESULTS

### Unbiased identification of LASV and LCMV NP interacting proteins in human cells

To identify novel host proteins that could take part in protein-protein interactions with the arenavirus NP, we applied an unbiased proteomics approach. We used a human lung epithelial cell line, A549 that was transfected with plasmids expressing either LCMV or LASV NPs fused to HA tag (NP-HA). As negative controls we included cells transfected with a plasmid encoding an HA-tagged protein unrelated to the arenavirus NP but with a similar molecular weight, the ubiquitin carboxyl-terminal hydrolase 14 (HA-USP14), as well as cells infected with a newly generated recombinant tri-segmented LCMV, expressing HA-tagged GFP (3rLCMV-HA-GFP, Fig. S1A). Cell lysates were immunoprecipitated with anti-HA monoclonal antibody (Fig. S1B) followed by mass spectrometry to identify interacting peptides, using a criteria of at least 2 unique tryptic peptides with a degree of confidence of 99% to identify each hit. This approach revealed a number of NP-interacting host proteins detected with LCMV, LASV, or both, NP-HA samples but never found in negative controls (Table 1). Thus, after conducting an unbiased proteomics approach in human cells, we pinpointed a selected number of host proteins as novel candidates involved in protein-protein interactions with LCMV and/or LASV NP.

**Table 1.** Host proteins interacting with the NP of LCMV, LASV, or both.

| GI | Annotation | LCMV | LASV | Total | NSC |
| --- | --- | --- | --- | --- | --- |
| **41399285** | **Chaperonin (HSPD1)** | 3 | 3 | 6 | 0.03892668 |
| 55956788 | Nucleolin # | 3 | 1 | 4 | 0.011823951 |
| 4506901 | Splicing factor, arginine/serine-rich 3 * | 3 | 1 | 4 | 0.004537574 |
| 7705813 | Ribosomal protein L26-like 1 | 2 | 1 | 3 | 0.01543909 |
| 4506633 | Ribosomal protein L31 isoform 1 | 1 | 2 | 3 | 0.012745707 |
| 4506619 | Ribosomal protein L24 | 2 | 1 | 3 | 0.010694274 |
| 4506623 | Ribosomal protein L27 | 2 | 1 | 3 | 0.008230397 |
| 15431297 | Ribosomal protein L13 | 1 | 2 | 3 | 0.007550774 |
| 4506607 | Ribosomal protein L18 | 2 | 1 | 3 | 0.005953904 |
| 4506675 | Ribophorin I precursor * | 2 | 1 | 3 | 0.001225967 |
| 57863257 | T-complex protein 1 isoform a | 0 | 2 | 2 | 0.020058442 |
| 57013276 | Tubulin, alpha, ubiquitous # | 1 | 1 | 2 | 0.019521338 |
| 4502643 | Chaperonin containing TCP1, subunit 6A isoform a | 0 | 2 | 2 | 0.01500201 |
| 24234688 | Heat shock 70kda protein 9 precursor * | 2 | 0 | 2 | 0.014046834 |
| 4506671 | Ribosomal protein P2 | 1 | 1 | 2 | 0.012759599 |
| 48762932 | Chaperonin containing TCP1, subunit 8 (theta) | 0 | 2 | 2 | 0.012183282 |
| 63162572 | Chaperonin containing TCP1, subunit 3 isoform a | 0 | 2 | 2 | 0.011693309 |
| 4506703 | Ribosomal protein S24 isoform c | 1 | 1 | 2 | 0.011032736 |
| 4506701 | Ribosomal protein S23 | 1 | 1 | 2 | 0.010261216 |
| 4504351 | Delta globin # | 2 | 0 | 2 | 0.010124655 |
| 5453603 | Chaperonin containing TCP1, subunit 2 | 0 | 2 | 2 | 0.008933907 |
| 38455427 | Chaperonin containing TCP1, subunit 4 (delta) | 0 | 2 | 2 | 0.008867607 |
| 4506741 | Ribosomal protein S7 | 1 | 1 | 2 | 0.00756368 |
| 4506667 | Ribosomal protein P0 | 1 | 1 | 2 | 0.00526533 |
| 117190254 | Heterogeneous nuclear ribonucleoprotein C isoform b | 1 | 1 | 2 | 0.005008034 |
| **156071459** | **Solute carrier family 25, member 5 (SLC25A5)** | 1 | 1 | 2 | 0.004924006 |
| **4758138** | **DEAD (Asp-Glu-Ala-Asp) box polypeptide 5 (DDX5)** | 1 | 1 | 2 | 0.004779654 |
| 5123454 | Heat shock 70kda protein 1A * | 2 | 0 | 2 | 0.004578327 |
| 4506725 | Ribosomal protein S4, X-linked X isoform | 1 | 1 | 2 | 0.00423095 |
| 45238849 | Poly (A) binding protein, cytoplasmic 3 * | 2 | 0 | 2 | 0.003538014 |
| **4557871** | **Transferrin (TF)** | 2 | 0 | 2 | 0.00213227 |
| **4885661** | **Viral oncogene yes-1 homolog 1 (YES1)** | 2 | 0 | 2 | 0.001370464 |
| **16751921** | **Dermcidin preproprotein (DC)** | 1 | 0 | 1 | 0.119585846 |
| 4759160 | Small nuclear ribonucleoprotein polypeptide D3 | 1 | 0 | 1 | 0.017718147 |
| **4504347** | **Alpha 1 globin (HGB)** | 1 | 0 | 1 | 0.015721736 |
| 5031839 | Keratin 6A * | 1 | 0 | 1 | 0.013994088 |
| 24307939 | Chaperonin containing TCP1, subunit 5 | 0 | 1 | 1 | 0.012340921 |
| 5453607 | Chaperonin containing TCP1, subunit 7 isoform a | 0 | 1 | 1 | 0.011736379 |
| 119703753 | Keratin 6B * | 1 | 0 | 1 | 0.010915491 |
| 15431288 | Ribosomal protein l10a | 1 | 0 | 1 | 0.010316443 |
| 58331185 | Chaperonin containing TCP1, subunit 7 isoform b | 0 | 1 | 1 | 0.009847254 |
| 4506743 | Ribosomal protein S8 | 0 | 1 | 1 | 0.008024566 |
| 4503471 | Eukaryotic translation elongation factor 1 alpha 1 # | 1 | 0 | 1 | 0.006352181 |
| 5902102 | Small nuclear ribonucleoprotein D1 polypeptide 16kda | 1 | 0 | 1 | 0.006253464 |
| 4759156 | Small nuclear ribonucleoprotein polypeptide A | 1 | 0 | 1 | 0.005277746 |
| **63252906** | **Tropomyosin 1 alpha chain isoform 7 (TPM)** | 1 | 0 | 1 | 0.005240579 |
| 17105394 | Ribosomal protein l23a | 1 | 0 | 1 | 0.004703057 |
| 66912162 | Histone cluster 2, h2bf # | 1 | 0 | 1 | 0.004441802 |
| 4506685 | Ribosomal protein S13 | 0 | 1 | 1 | 0.003684569 |
| 51477708 | Heterogeneous nuclear ribonucleoprotein D isoform d | 1 | 0 | 1 | 0.003543757 |
| 5174431 | Ribosomal protein L10 | 1 | 0 | 1 | 0.003428397 |
| 47271443 | Splicing factor, arginine/serine-rich 2 # | 1 | 0 | 1 | 0.00336725 |
| 76496472 | Ribosomal protein L3 isoform b | 0 | 1 | 1 | 0.003143333 |
| 57863259 | T-complex protein 1 isoform b | 1 | 0 | 1 | 0.001829618 |
| 55770868 | Tubulin, beta polypeptide 4, member Q # | 1 | 0 | 1 | 0.0016905 |
| **70906435** | **Fibrinogen, beta chain preproprotein (FG)** | 1 | 0 | 1 | 0.001515605 |
| 15431310 | Keratin 14 | 1 | 0 | 1 | 0.001185735 |
| **87196351** | **DEAD/H (Asp-Glu-Ala-Asp/His) box polypeptide 3 (DDX3)** | 1 | 0 | 1 | 0.000845418 |
| **110431348** | **Deleted in colorectal carcinoma (DCC)** | 1 | 0 | 1 | 0.000514279 |

Hits were considered positive when 2 unique tryptic peptides were detected in either LCMV or LASV samples and never (or only 1 tryptic peptide\*) in negative controls (HA-USP14 or 3rLCMVGFp-HA). Samples belong to 4 independent experiments. Hits were first ranked in groups according to the number of times detected (6 to 1) in the 8 samples (4 LCMV + 4 LASV). Within each group, hits were ranked according to the highest NSC (Normalized spectral counts) value detected for each hit. #Hits detected in THP1 negative controls from unrelated MS studies. **Bold**: Hits selected for siRNA functional screening. GI: Gene identification (NCBI databank).

### Functional screening with NP interacting candidates singled out DDX3

To functionally characterize the role of newly identified NP-interacting candidates in arenavirus infection, we conducted a loss-of-function assay to monitor viral growth in cells treated with small interfering RNAs (siRNAs) directed against selected NP interacting candidates (Table 1, bold). To select these candidates, hits in Table 1 were prioritized by further filtering out proteins for which only one unique tryptic peptide was detected in negative control samples, proteins that were detected in THP1 negative control cells from previous unrelated mass spectrometry studies, as well as arbitrarily excluding ribosomal and ribonucleoproteins. A549 cells were incubated with targeting siRNA or scrambled siRNA (Scr1-siRNA or Scr2-siRNA) prior to LCMV infection. Approximately 90% of the cells incorporated siRNA oligonucleotides (Fig. S1C) and cell viability was comparable for all siRNAs tested (Fig. S1D, representative result is shown). Cells transfected with DDX3-specific, but not scrambled, siRNA showed both reduced levels of DDX3 protein (Fig. S1E) and a reduction of LCMV titers (i.e. ~0.5 log) (Fig. 1A). In contrast, viral yields were unaffected upon transfection with siRNAs against other candidates. Note that a negative result in this screen could have resulted from a non-essential role of the target gene and/or insufficient down-regulation of the target gene in the conditions used in our assay. Overall, these results singled-out DDX3 as a host factor that could potentially play a pro-viral role in arenavirus life cycle.

**Figure 1.**
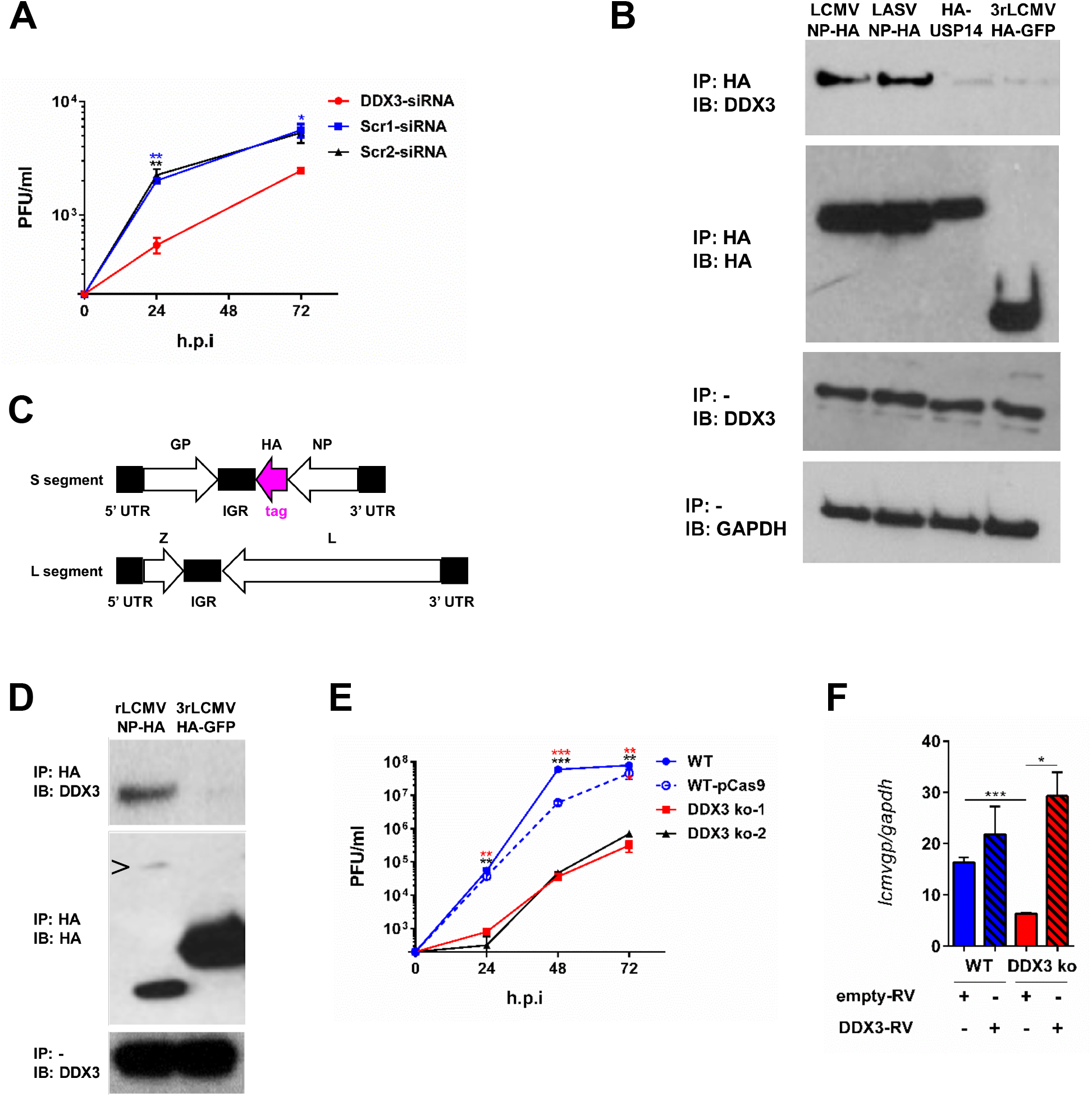
DDX3 interacted with LASV and LCMV NPs and promoted LCMV growth in human cells. **A.** A549 cells were transfected for 60h with targeting siRNAs specific for DDX3 or scrambled siRNA controls followed by infection with LCMV C113 (M.O.I. 0.005). Viral titers in cell culture supernatants harvested 24 and 72 h.p.i. are shown. B. A549 cells were transfected for 24h with plasmids encoding LCMV-NP-HA, LASV-NP-HA or HA-USP14, or infected with 3rLCMV-HA-GFP, lysed and immunoprecipitated with anti-HA agarose beads (IP HA); eluates were analyzed by Immunoblot (IB). Immunoblots with anti-DDX3 and anti-GAPDH (load control) Abs were performed in input samples. **C.** Schematics of the genome of rLCMV-NP-HA. White: ORFs of viral proteins. Pink: HA tag. Black: viral untranslated regions. **D.** A549 cells were infected with rLCMV-NP-HA or 3rLCMV-HA-GFP for 24h, lysed in buffer containing RNAseA 0.1 mg/ml, immunoprecipitated (IP) and analyzed by Immunoblot (IB). > indicates NP-HA band. **E.** Viral titers in supernatants from DDX3 ko-1, DDX3 ko-2, WT-pCas9 (control) and WT A549 cells infected with LCMV Cl13 (M.O.I. 0.5) were quantified at 24, 48 and 72 h.p.i. **F.** DDX3 ko-1 and WT A549 cells transduced with RV expressing DDX3 or empty-RV were infected with LCMV C113 (M.O.I. 0.5) for 24h and viral RNA levels *(Icmvgp)* were determined relative to *gapdh* by RT-qPCR. Data are representative of 2 (**A**, **D** and **F**) or 4 (**B** and **E**) independent experiments. * p<0.05, ** p<0.01, ***p<0.001. Stars colors represent: DDX3 vs. Scr1(blue) or Scr2 (black) (A) and WT A549 vs DDX3 ko-1 (red) or vs DDX3 ko-2 (black) (**E**).

### DDX3 associated to LCMV NP under both transfection and infection conditions

We next attempted to validate arenavirus NP interaction with DDX3. For that we transfected cells with plasmids encoding LCMV or LASV NPs, and as negative controls we used cells transfected with HA-USP14 or infected with 3rLCMV-HA-GFP. Cell lysates were immunoprecipitated with an anti-HA mAb, and analyzed by immunoblotting with an anti-DDX3 Ab. DDX3 levels were higher in immunoprecipates from cells transfected with LCMV or LASV NP-HA than in the negative control samples (Fig. 1B, top panel). Similar levels of DDX3 and GAPDH were detected in input lysates from all samples (Fig. 1B, middle and bottom panels). These results validated that DDX3 interacted with both LCMV and LASV NPs in transfected cells. To test whether NP and DDX3 interact in the context of arenavirus infection, we generated a recombinant LCMV expressing a HA-tagged version of NP (Fig. 1C). Viral titers of rLCMV-NP-HA were typically ~1 log lower than those obtained with WT rLCMV, but no differences in size or shape of plaques were observed and importantly both viruses replicated to similar titers *in vivo*, revealing no gross changes in viral fitness (Fig. S1F). DDX3 was immunoprecipitated from cells infected with rLCMV-NP-HA but not from control samples infected with 3rLCMV-HA-GFP (Fig 1D top panel). Partially cleaved NP-HA and GFP-HA in immunoprecipitates and DDX3 levels in input samples were readily detectable (Fig. 1D, middle and bottom panels). Together these results demonstrated that DDX3 associated with OW arenavirus NP both in transfected and infected cells.

### DDX3 promoted LCMV and LASV growth in human cells

To further investigate the role of DDX3 in OW arenavirus infection we generated two DDX3 knockout (ko) cell lines by using different non-overlapping RNA guides and CRISPR/Cas9 gene editing, and processed in parallel a control cell line transfected with a plasmid lacking RNA-guides (WT-pCas9). Immunoblot of cell lysates showed that DDX3 was undetectable in both ko cell lines (Fig. S2A). To investigate the impact of DDX3 gene deletion on LCMV viral growth, the DDX3 ko cell lines, WT and WT-pCas9 control cells were infected with LCMV and viral RNA synthesis and production of infectious progeny were monitored over time. Cell viability at the time of infection was similar for all cell lines (Fig. S2B). Instead, production of LCMV infectious progeny was largely reduced (i.e. ~ 2 *log*) at all times examined, which correlated with reduced levels of viral RNA in DDX3 ko cells at 8 and 24 hpi (Fig 1E & S2C). Conversely, when DDX3 ko cells were infected with Sendai virus (SeV) we did not observe any reduction in viral RNA (Fig. S2D). Importantly, reconstitution of DDX3 protein expression in DDX3 ko cells transduced with a retrovirus (RV) encoding DDX3 (Fig. S2E) resulted in significantly increased viral RNA levels compared to DDX3 ko cells transduced with empty RV (Fig. 1F). Overexpression of DDX3 in WT cells transduced with DDX3-RV did not, however, cause any significant changes in viral RNA amounts respect to WT cells transduced with empty RV (Fig 1F, blue bars). These data supported that reduced LCMV growth in DDX3 ko cells was due to lack of DDX3 expression rather than off-target effects.

We next evaluated the role of DDX3 during infection with the HF OW arenavirus LASV. For that, we performed similar infection experiments in WT versus DDX3 ko cell lines in BSL-4. Quantification of infected cells via confocal microscopy revealed a significant reduction in LASV growth in both DDX3 ko cell lines compared to WT controls at all the M.O.I. tested (Fig. 2A&B). Consistently, we detected 100 to 1000-fold less LASV RNA in the culture supernatants of both DDX3 ko versus WT cell lines (Fig. 2C). LASV infection rates were increased in both DDX3 ko cell lines when DDX3 expression was reconstituted via RV transduction, reaching statistical significance at M.O.I 0.05 and 0.1 (Fig. 2D).

**Figure 2.**
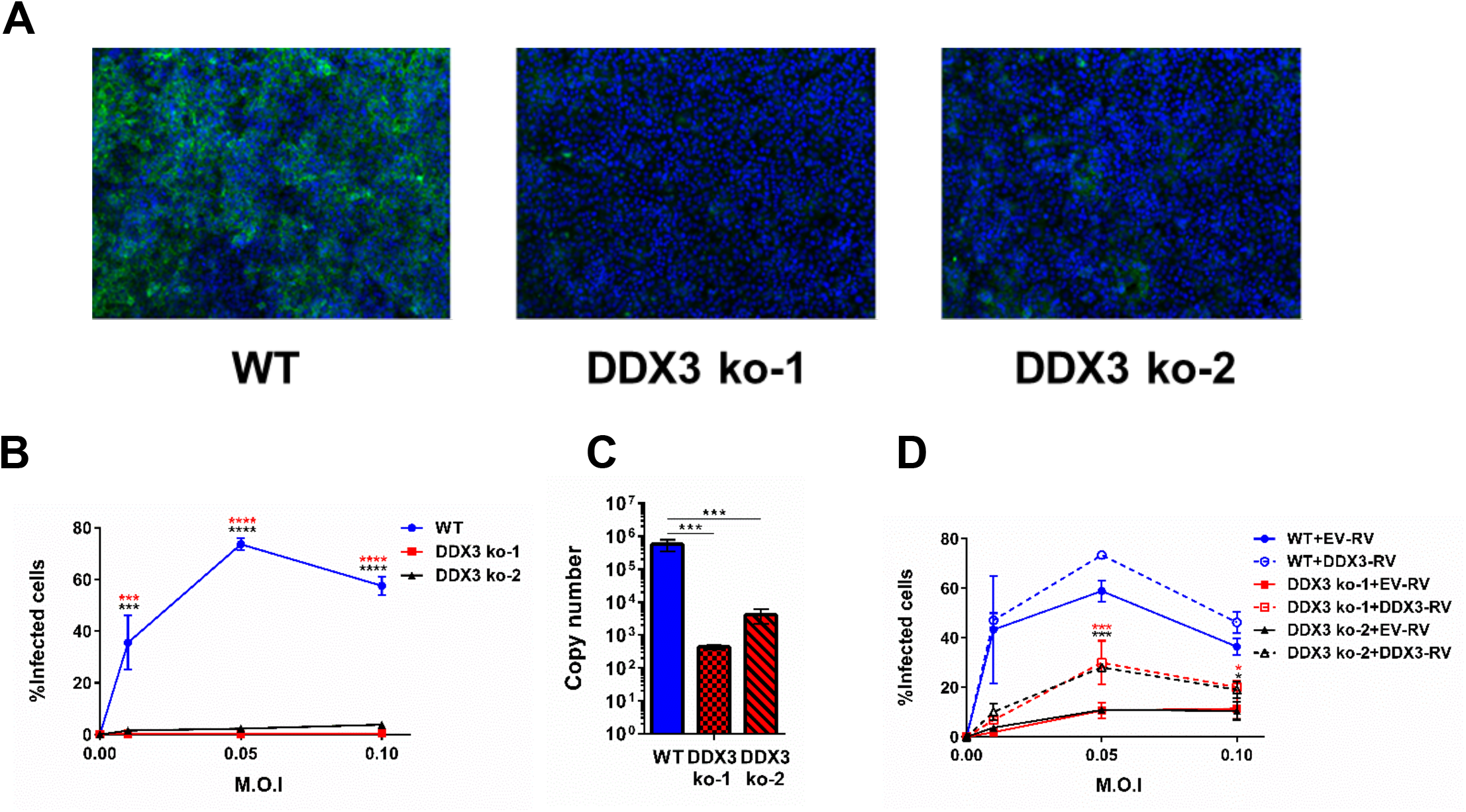
DDX3 promoted LASV growth in human cells. **A** to **C.** DDX3 ko-1, DDX3 ko-2 and WT A549 cells were infected with LASV (strain Josiah) at the indicated M.O.I. for 48h. **A.** Cells were fixed with 10% formalin for 72h and stained with anti-GP antibodies and Hoechst, for confocal microscopy. Representative images are shown. **B.** Number of infected cells were calculated by high-content quantitative image-based analysis. **C.** Viral RNA yields in tissue culture supernatants were determined by qRT-PCR. **D.** DDX3 ko-1, DDX3 ko-2 and WT A549 cells were transduced with empty-RV (EV-RV) or RV encoding DDX3 (DDX3-RV) prior to LASV infection and then processed as in **B.** All data are representative of 2 independent experiments. * p<0.05, ** p<0.01, ***p<0.001. Star colors represent: WT-A549 vs. DDX3 ko-1 (red) or vs. DDX3 ko-2 (black) (B) and DDX3 ko-1+EV-RV vs DDX3 ko-1+DDX3-RV (red) or DDX3 ko-2+EV-RV vs DDX3 ko-2+DDX3-RV (black).

Altogether, these observations provided strong evidence that DDX3 was a pro-viral host factor, which promoted optimal viral growth during infection with OW arenaviruses LCMV and LASV in human cells.

### DDX3 contributed to the IFN-I suppression observed upon arenavirus infection

DDX3 was previously shown to interact with several components of the IFN-I pathway and to enhance IFN-I production [24–27]. To investigate a putative role for DDX3 in IFN-I induction after arenavirus infection we quantified *IFNB* transcript levels in WT and DDX3 ko cells after LCMV infection. Consistent with the potent capacity of arenaviruses to suppress IFN-I induction [15,16,28–30], *IFNB* was undetectable in WT cells infected with LCMV (Fig. 3A, blue line). In contrast, increasing amounts of *IFNB* transcript were detected in DDX3ko cells during the first 24 hours after LCMV infection (Fig. 3A, red line). While this effect was more profound at M.O.I 0.5, it was also significant at M.O.I 0.1 and 2.5 (Fig. S3A) and it was attenuated when DDX3 levels were reconstituted in DDX3 ko cells, at both 24 and 48 h.p.i (Fig 3B). Similar results were obtained when IFN-I bioactivity was quantified in the culture supernatant at 48 h.p.i (Fig. 3C). These findings are in sharp contrast to the previously reported role of DDX3 in promoting IFN-I induction [24,25] and revealed for the first time a suppressive role of DDX3 on *IFNB* transcription, suggesting that the influence of DDX3 on IFN-I production is context dependent.

**Figure 3.**
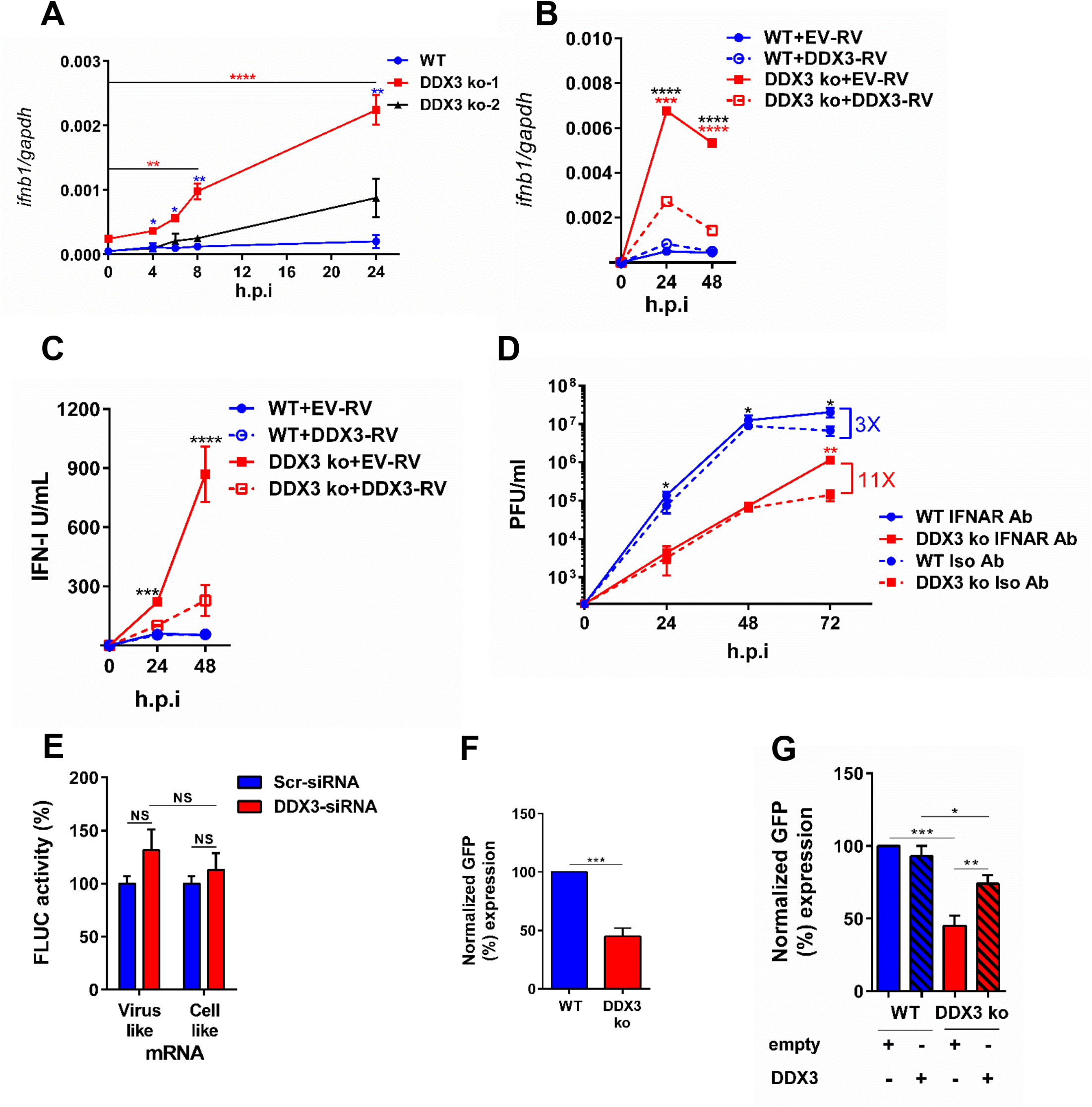
DDX3 contributed to IFN-I suppression and viral replication/transcription after arenavirus infection in human cells. **A.** DDX3 ko-1, DDX3 ko-2 and WT A549 cells were infected with LCMV C113 (M.O.I. 0.5) for the indicated times. Total RNA in cell lysates was extracted and normalized viral RNA levels (*ifnb/gapdh*) determined by qRT-PCR. B-C. DDX3 ko-1 and WT A549 cells were transduced with empty-RV (EV-RV) or RV encoding DDX3 (DDX3-RV) before infection, and processed as in **A** for quantification of *ifnb/gapdh* transcripts via qRT-PCR (**B**) or determination of bioactive IFN-I levels in cell culture supernatants at indicated hp.i. (**C**). **D.** DDX3 ko-1 and WT A549 cells were pre-incubated for 2 h and infected with LCMV Cl13 (M.O.I. 0.5) in the presence of anti-IFNAR mAb (IFNAR Ab) or Isotype control (Iso Ab), which were left for the remaining of the culture. Viral titers were determined at indicated h.p.i. E. Translation assay performed in HEK-293T cells treated with DDX3 or Scr siRNA. **F.** Minireplicon assay performed in DDX3 ko-1 or WT A549 cells **G.** DDX3 ko-1 or WT cells were transfected with 0.4 μg of empty plasmid or plasmid expressing DDX3 and used for minireplicon assay. 100% value was given to WT A549 transfected with empty plasmid. Data are representative of 2 (**A-D** & **F-G**) or 3 (**E**) independent experiments. * p<0.05, ** p<0.01, ***p<0.001. Stars represent: DDX3ko vs WT (blue) or vs DDX3 ko at the indicated h.p.i. versus time=0 (red) (**A**), DDX3 ko-EV vs WT-EV (black) or vs DDX3 ko-RVDDX3 (red) (**B**), WT-EV vs DDX3 ko-EV (**C**) and DDX3 ko-IFNAR vs WT-IFNAR (black) or vs DDX3 ko-Isotype (red) (**D**).

To test whether increased IFN-I levels could contribute to the diminished LCMV growth in the absence of DDX3, we incubated WT and DDX3 ko cells with anti-IFN-I receptor (IFNAR) mAb or isotype control before and throughout LCMV infection. We observed an 11-fold increase in LCMV titers at 72 (but not 24 or 48) h.p.i. in DDX3 ko cells incubated with anti-IFNAR versus isotype mAb, in contrast to a 3-fold increase when anti-IFNAR was blocked in WT cells (Fig. 3D). These results suggested that while the pro-viral effect of DDX3 was partially due to IFN-I suppression at late time points after infection, DDX3 pro-viral activity was independent of IFN-I signaling at early time points. This was consistent with significant reduction of LCMV RNA levels following DDX3 depletion via DDX3-specific (compared to scrambled) siRNA in Vero cells, which naturally lack the IFN-I system [31] (Fig. S3B&C).

Altogether these results pointed to a IFN-I independent, early mechanism involved in DDX3 enhancement of arenavirus propagation, and also revealed a previously unrecognized role of DDX3 as a suppressor of *IFNB* transcription, partially explaining DDX3 pro-viral role late after LCMV infection.

### DDX3 promoted arenavirus replication/transcription but was dispensable for translation

Given that DDX3 is able to promote translation of both viral and cellular mRNAs [32,33], we next investigated whether DDX3 also played a role in arenavirus mRNA translation. For that we used a recently reported arenavirus translation assay based on capped synthetic RNAs carrying the reporter firefly Luciferase (FLUC) open reading frame [34]. Quantification of the luciferase reporter activity after transfection of a Tacaribe (TCRV) mRNA analog, or a cell-like transcript as control, in cells treated with DDX3-specific indicated that the translation of both the viral and cellular mRNA analogs was unchanged compared to control cells transfected with scrambled siRNA (Fig. 3E and S4). These results suggested that the reduction in viral growth observed in DDX3 deficient cells was unlikely due to reduced RNA translation. Thus, we next investigated DDX3 role in arenavirus replication/transcription by using a well-established LCMV minireplicon system that assesses the activity of the intracellularly reconstituted viral ribonucleoprotein (vRNP) responsible for directing viral RNA replication and gene transcription [12]. We used a LCMV S segment-based minireplicon where the *Gaussia* luciferase (Gluc) and GFP reporter genes substituted for NP and GPC genes, respectively, within the S genome RNA (MG/Gluc-GFP). Levels of GFP expression in cells transfected with MG/Gluc-GFP together with plasmids expressing the viral trans-acting factors NP and L polymerase, serve as surrogate of the vRNP activity. GFP signal showed a 60% drop in minireplicon activity in DDX3 ko cells compared to WT controls (Fig. 3F), which was significantly increased when DDX3 ko cells were transfected with a DDX3-encoding, but not empty, plasmid (Fig. 3G). These results suggested that the pro-viral role of DDX3 on arenavirus multiplication was likely related to a novel function of DDX3 in promoting arenavirus replication and/or transcription.

### DDX3 interacted with New World arenavirus NPs and promoted JUNV growth in human cells

We next investigated whether DDX3-NP interaction was also conserved in NW arenaviruses. For that, we transfected cells with plasmids expressing the NP of JUNV, MACV or TCRV tagged with HA and prepared cell lysates that were examined by immunoprecipitation with anti-HA mAb followed by Immunoblot with anti-DDX3 mAb. DDX3 levels were significantly enriched in immunoprecipitates from cells transfected with plasmid encoding NW arenavirus HA-tagged NPs compared to cells transfected with empty plasmids (Fig 4A), indicating that DDX3 interacted with the NP of all the NW arenaviruses tested. We next investigated whether DDX3 also played a pro-viral role in NW arenavirus growth. For this we first infected the two DDX3 ko cell lines (Fig. S2A) with the vaccine strain of JUNV (Candid#1) [7]. We observed that the number of cells infected with Candid#1 was dramatically reduced to near-undetectable levels in both DDX3 ko cell lines versus WT controls, an effect that was maintained at all M.O.I tested (Fig. 4B). Importantly, DDX3 reconstitution resulted in a significant increase in infection rate, which reached statistical significance at M.O.I 1 (Fig. 4C). To confirm these results with a highly pathogenic strain of JUNV, both DDX3 ko cell lines were infected with JUNV Romero strain in BSL-4. Quantification of infected cells and JUNV RNA levels in cell culture supernatants indicated a dramatic decrease in both parameters in DDX3 ko versus WT control cells, regardless of the M.O.I used (Fig. 4D-F). This effect was partly reverted in both DDX3 ko cell lines when DDX3 expression was reconstituted, reaching statistical significance at all M.O.I tested (Fig. 4G). These results indicated that, as with OW arenaviruses, the NP of NW arenaviruses interacted with DDX3 and that this helicase was required for optimal growth of JUNV in human cells.

**Figure 4.**
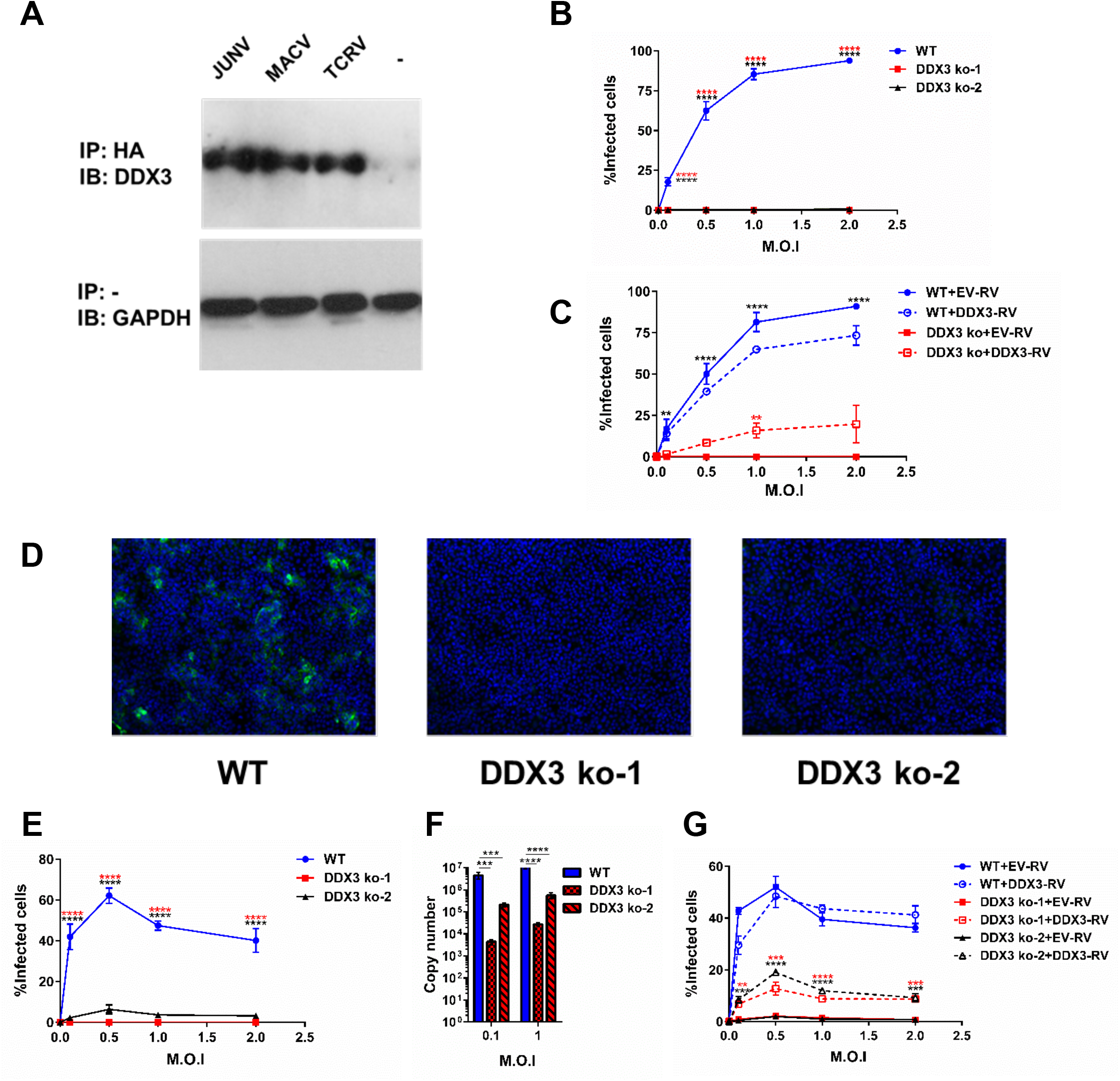
DDX3 interacted with New World arenavirus NP and promoted JUNV growth in human cells. **A.** A549 cells were transfected with plasmid encoding JUNV, MACV or TCRV NP-HA for 24h, lysed and immunoprecipitated with anti-HA agarose beads (IP HA). Eluates (middle panel) or input samples (load control, lower panel) were analyzed by Immunoblotting (IB) with anti-DDX3 or anti-GAPDH, respectively. **B-G**. DDX3 ko-1, DDX3 ko-2 and WT A549 cells were infected with JUNV Candid 1 (**B-D**) or Romero (**E-G**) strains at the indicated M.O.I. for 24h. Cells were fixed with 10% formalin for 72h and stained with anti-GP antibodies and Hoechst, for confocal microscopy. Representative images for infected cells are shown (**D**). Number of infected cells was determined by high-content quantitative image-based analysis (**B**, **C**, **E** and **G**). Viral RNA yields in tissue culture supernatants were determined by qRT-PCR (**F**). When indicated, (**C** and **G**) DDX3 ko-1 and WT A549 cells were transduced with empty-RV (EV-RV) or RV encoding DDX3 (DDX3-RV) before infection, and processed as in B and E, respectively. Data are representative of 2 (A-F) or 1(G) independent experiments. ** p<0.01, ***p<0.001. Star colors represent: DDX3 ko-EV vs WT-EV (black) or vs DDX3 ko-RVDDX3 (red) (**C**), WT vs DDX3ko-1(red) or vs DDX3ko-2 (black) (**E**) DDX3 ko-1+EV-RV vs DDX3 ko-1+DDX3-RV (red) or DDX3 ko-2+EV-RV vs DDX3 ko-2+DDX3-RV (black) (**G**).

## DISCUSSION

Arenaviruses are endemic in their natural rodent hosts and often infect humans, with LASV causing thousands of lethal hemorrhagic fever cases each year [1–3]. Moreover, these viruses have been listed among the top priority emerging pathogens that are likely to cause a severe outbreak in the near future [10]. Currently there is no FDA-approved vaccine against arenavirus infections and there are limited therapeutic options that include Ribavirin, a compound with many side effects that requires administration in the first days of infection to show partial effectiveness [9]. Thus, there is an urgent need to develop new strategies to treat or prevent arenavirus infection in humans. A better understanding of arenavirus host interactions will not only inform about fundamental cellular processes exploited or subverted by these viruses, but could also help identify such intervention strategies. Given that therapeutic targeting of host (rather than viral) factors would minimize arenavirus escape mutants, we used an unbiased approach to identify arenavirus interacting candidates in human cells. Because of its abundant expression and key role in viral fitness and immune-evasion, we focused on host factors that interact with the viral protein NP. Among several newly identified NP interacting candidates, we established DDX3, an ATP-dependent RNA helicase, as an arenavirus target that is exploited to suppress host immunity and promote viral replication/transcription.

Notably, we have biochemically validated NP-DDX3 interaction in both NP-transfected and virus-infected cells, for both OW and NW arenavirus NPs. These results are consistent with a very recent report that examined the interactome of LCMV and JUNV NPs via mass spectrometry and documented (but did not biochemically validate or functionally characterize) DDX3 among numerous other NP interacting candidates [23]. While some of the NP-interacting proteins reported by King et al. were also detected in our NP interactome (e.g. DDX3, DDX5, SCLC25A5, HSPD-1 and TPM), other candidates were only detected by one of the two studies. These discrepancies could be related to the use of LCMV-infected cells [23] versus NP-transfected cells in our initial mass spectrometry approach, by the lower sensitivity of our method (i.e. lower spectral count values) and/or by the more stringent cut-off criteria that we used based on 99% confidence for peptide detection and absence of hits in two negative controls. Interestingly, neither we, nor King et al. [23] detected some previously documented NP-interacting host proteins including IKKε, MDA5 and RIG-I [20,21]. Most importantly, our functional studies demonstrated that DDX3 was critical for optimal arenavirus multiplication, as DDX3 inhibition via either siRNA or CRISPR/Cas9 gene editing led to a significant reduction in viral titers after infection not only with LCMV, but also with the HF arenaviruses LASV and JUNV. Thus, although targeting DDX3 should be carefully weighed in the context of its many physiological roles [35–38], our results raised the attractive possibility that treatment with DDX3 inhibitors could be a viable and broadly-effective approach to curtail viral replication and alleviate arenavirus infections in humans, either alone or in combination with ribavirin. Interestingly, DDX3 appears to represent a convergent viral target, as it has been reported to interact with multiple types of viral proteins [24,39–42]. Although the outcome of these interactions has revealed both pro-viral and antiviral effects of DDX3, the fact that DDX3 is targeted by distantly related viruses, suggests an important role for DDX3 in antiviral defense.

In other infections a pro-viral role of DDX3 has been related to distinct mechanisms [43]. While during Japanese encephalitis virus and norovirus infections DDX3 helicase activity is crucial for viral replication via an unknown mechanism [44,45], upon West Nile virus infection DDX3 is known to be sequestered from stress granules and processing bodies towards viral replication sites [46]. As for human immunodeficiency virus (HIV) infection DDX3 is important for both the nuclear export of unspliced vRNA and the translation of HIV transcripts [41,47]. Here, we showed that DDX3 deletion resulted in inhibition of arenavirus multiplication that correlated with a decreased in vRNP activity as determined in a cell-based LCMV minireplicon assay. Because DDX3 is involved in dsRNA unwinding via its helicase domain [48], it is possible that NP interacts with DDX3 to recruit into the virus replication complex a helicase activity required to facilitate RNA synthesis by the arenavirus polymerase. Alternatively, given that DDX3 is critical for stress granule (SG) formation independently of its helicase domain [49], it is also possible that arenavirus NP interaction with DDX3 has evolved to recruit SG (and other) proteins into the replication transcription complexes (RTC) organelles [50], thereby facilitating viral replication.

DDX3 has been reported to exert antiviral activity by enhancing IFN-I production [51,52] and to up-regulate IFN-I via interaction with IKKε or TBK1 [24,25,27], as well as to interact with RIG-I and MDA5 that bind to MAVS and facilitates IFNβ induction [52]. DDX3 can also act as a transcriptional regulator by interacting with IFNβ promoter [25]. Importantly, enhancement of IFN-β production by DDX3 has been proposed to mediate its antiviral activity against vesicular stomatitis virus [52]. Very recently, DDX3 has also been reported to initiate MAVS signaling by sensing abortive RNA in HIV infected dendritic cells [51]. Strikingly, our experiments revealed a novel and unexpected role for DDX3 in suppressing *IFNB* transcription upon LCMV infection. Such IFN-I suppression appeared to partially contribute to DDX3 pro-viral activity late in infection, an effect that is expected to be amplified *in vivo* considering that studies in IFN-AR deficient animal models demonstrated critical roles played by IFN-I in promoting activation of almost all immune cells [53] and protection against arenavirus multiplication [54–56]. Although, IFNAR blockade has been shown to relieve immunosuppression during chronic LCMV infection in mice, it also enhanced viral titers initially [57,58]. The last-mentioned result is also consistent with evidence that treatment with rIFNα/β early after LCMV infection promotes viral clearance [59], supporting a beneficial effect of enhancing IFN-I during arenavirus infections *in vivo*. It is possible that NP-sequestration of DDX3 may carry over some of the known DDX3-interacting proteins that participate in IFN-I induction (i.e. IKKε, TBK1, RIGI, MDA5 and/or MAVS) [22,53], counteracting the formation of macromolecular complexes required for IFN-I synthesis. Furthermore, the possible requirement of DDX3 in the formation of RTC organelles may diminish IFN-I induction by enabling the compartmentalization of viral RNA replication and transcription; therefore limiting recognition of dsRNA intermediates by innate sensing receptors. In addition, given that arenavirus mRNAs are not polyadenylated [60] and are therefore expected to be sensed by DDX3 [51], it is possible that arenavirus NP-DDX3 interaction has evolved to sequester DDX3 into compartments that are devoid of viral mRNAs [50], thereby preventing their recognition. Thus, although further studies are necessary to fully understand the underlying mechanisms, it is remarkable that deletion of a single host protein (i.e. DDX3) resulted in (at least partial) counteraction of the long-evolved capacity of arenavirus to suppress *IFNB* induction [15,16,30].

Our findings using an unbiased proteomic approach followed by biochemical validation in infected cells, identified DDX3 as a novel interacting partner of OW and NW arenavirus NP. Importantly, we have also uncovered two previously unrecognized DDX3-dependent strategies by which arenaviruses might counteract the host cell IFN-I response and exploit the host cellular machinery to maximize their multiplication. These findings provide the fundamental knowledge to consider DDX3 inhibitors as a potential therapeutic approach to treat infections by human pathogenic arenaviruses.

## MATERIAL AND METHODS

### Cells

A549 (Human lung epithelial cells, ATCC^®^ CCL-185^TM^, Manassas, VA), BHK-21 (Newborn Hamster kidney fibroblast cells, ATCC^®^ CCL-10^TM^) and HEK-293T (Human epithelial kidney cells, ATCC^®^ CRL-11268^TM^) were cultured in Dulbecco’s Modified Eagle Medium (DMEM) (11965-118, Gibco, Grand Island, NY, USA) supplemented with 2 mM L-glutamine (25030081, Thermo Scientific), 50 U/mL penicillin-streptomycin (15140-163, Gibco), plus 10% heat-inactivated FBS (Lonza). BHK-21 were also supplemented with 20% Tryptose Phosphate Broth (18050039, Thermo Scientific). HEK-293T cells were supplemented with sodium pyruvate (1 mM) and non-essential amino acids (0.1 mM). HEK-Blue^TM^ IFN-α/β cell line (InvivoGen) was maintained in HEK-293T media supplemented with 10μg/ml blasticidin (InvivoGen) and 200 μg/ml Zeocin (InvivoGen). Vero E6 cells *(Cercopithecus aethiops* kidney epithelial cells, ATCC^®^ CCL-81^TM^) were cultured in Minimum Eagle Medium (MEM) (11095-080, Gibco), supplemented with 2 mM L-glutamine, 50 U/mL penicillin-streptomycin and 7.5 % heat-inactivated FBS. A549, BHK-21, HEK-293T and Vero E6 cell lines were originally provided by J.C. de la Torre (The Scripps Research Institute, La Jolla, CA) and HEK-Blue^TM^ IFN-α/β cell line by S. Sharma (La Jolla Institute for Allergy and Immunology, La Jolla, CA)

### Viruses

LCMV Cl13 stocks were produced in BHK-21 cells and viral titers were determined by M6 well plaque assay on Vero cells. LCMV Cl13 infections were performed in BSL-2 facilities as previously described [61]. All work with highly pathogenic arenaviruses was performed at the United States Army Medical Research Institute of Infectious Diseases (USAMRIID) at Fort Detrick, Frederick, MD, USA, within maximum containment (BSL-4). JUNV Romero or Candid#1 strain and LASV Josiah strain viruses were propagated in Vero cells and viral infectivity was titrated by plaque assays as previously reported [62]. Sendai Virus infections were performed with Cantell strain.

### Generation of recombinant viruses

3rLCMV-HA-GFP was generated by modifying the previously described 3rLCMV-GFP virus [63] through the insertion of a HA-FLAG tag sequence (YPYDVPDYADYKDDDDK) in the N-terminal end of the GFP ORF (located in place of NP ORF) in one of the pol-I S vectors by multi-fragment assembly [64] using Phusion High Fidelity Polymerase (Thermofisher Scientific). For the viral rescue, BHK-21 cells (2 x 10^6^ cells per M6 well) were transfected for 5h by using 2μl of Lipofectamine 2000 (Invitrogen) per microgram of plasmid DNA. The plasmid mixture was composed of 0.8μg of pC-NP, 1μg of pC-L, 1.4μg of pol-I L and 0.8μg of each of the two pol-I S vectors. We confirmed expression of HA-GFP protein (~27kDA) in cells infected with 3rLCMV-HA-GFP by flow cytometry and Immunoblot with anti-HA Ab. Recombinant rLCMV-NP-HA was generated similarly, but using one single pol-I S vector expressing a modified NP ORF with the HA tag coding sequence on its C-terminal domain, as mentioned for 3rLCMV-HA-GFP. Primers for LCMV-HA-GFP: 1^st^ round (addition of FLAG seq): Fragment 1 (Fr1-Fw: CGGACATCT GGTCGACCTCCAGCATCG and Fr1-Rv: GATTACAAGGATGACGACGAT AAGT AA GACCCTCTGGGCCTCCCTGACTCTCCACCTCTTTCGAG) and Fragment 2 (Fr2-Fw: CTTATCGTCGTCATCCTTGTAATCCATCTTGTTGCTCAATGGTTTCTCAAGACAAATGCGCAATCAAATGC and Fr2-Rv: CGATGCTGGAGGTCGACCAGATGTCCG). 2^nd^ round (addition of HA seq): Fragment 3 (Fr1-Fw and Fr3-Rv: TACCCTTATGATGTCCCAGATTATGCCGATTACAAGGATGACGACGATAAGGTG AGC) and Fragment4 (Fr4-Fw: GGCATAATCTGGGACATCATAAGGGTACATCTTGTTGCTCAATGGTTTCTCAAGACAAATGCGCAATC and Fr2-Rv). Primers for rLCMV-NP-HA: Fragment 5 (Fr5-Fw: CCTACAGAAGGATGGGTCAGATTGTGACAATGTTTGAGGCTC and Fr5-Rv: TCCGGAGCCTACCCTTATGATGTCCCAGATTA TGCCTAAGACCCTCTGGGCCTCCCTGACTCTCCACCTCTTTCGAGGTGG, Fragment6 (Fr6-Fw: GGCATAATCTGGGACATCATAAGGGTAGGCTCCGGAGAGT GT CACAACATTT GGGCCT CT AAAAATT AGGT CAT GT GGCAG and Fr6-Rv: GGTTGGACTTCTCTGAGGTCAGCAATGTTCAG) and Fragment 7 (Fr7-Fw: CTGAACATTGCTGACCTCAGAGAAGTCCAACC and Fr7-Rv: GAGCCTCAAACA TTGTCACAATCTGACCCATCCTTCTGTAGG).

### Plasmids

pol-I S, pol-I L, pC-L, pC-NP, as well as pCAGGS plasmids encoding LASV, JUNV, MACV and TCRV NPs are described elsewhere [16]. HA-USP14: plasmid encoding ubiquitin-specific protease 14 fused to HA epitope to its N-terminal end [65]. Plasmid expressing DDX3 was constructed by inserting DDX3 cDNA in pCIneo-HA vector in EcoRI/NotI sites as previously described [66]. pSpCas9(BB)-2A-GFP construct is described in [67].

### Mice

C57BL/6 mice were purchased from The Jackson laboratory (Bar Harbor, ME). All mice were bred and maintained in a closed breeding facility and mouse handling conformed to the requirements of the National Institutes of Health and the Institutional Animal Care and Use Guidelines of UCSD. 6-8 weeks old mice were infected i.v with 5 x 10^6^ PFU of rLCMV or rLCMV-NP-HA.

### Antibodies

Anti-HA-Tag (C29F4) Rabbit mAb #3724 (dilution 1:3500), Anti-GAPDH (14C10) Rabbit mAb #2118 (1:5000), Anti-rabbit IgG, HRP-linked Antibody (1:5000) were obtained from Cell Signaling Technologies. Anti-DDX3 Rabbit Ab (1:5000, SAB3500206) was obtained from Sigma-Aldrich. Anti-IFNAR2 antibody (#21385-1, PBL Interferon Source) and Isotype control IgG2a antibody (#554126, BD Pharmingen Product) were both used at 5 μg/ml. Anti-LASV (L-52-161-6) and anti-JUNV (GD01) antibodies were obtained from the US Army research Institute of Infectious Diseases (USAMRIID) archives (PMID: 20686043, PMID: 22607481).

### Immunoprecipitation and Immunoblot

A549 cells (100,000 cells/ml) were plated on M12 wells. For Mass-spectrometry, four M12 plates were used for each experimental condition. Cells were transfected with 1μg of plasmid/well, encoding different arenavirus nucleoproteins fused to HA epitope in its C-terminal end (NP-HA [68] or plasmid encoding HA-USP14. Alternatively, cells were infected with 3rLCMV-HA-GFP with an m.o.i of 0.05. Media was replaced 6h later, and 24 h.p.t. cells were washed twice with PBS and lysed with 200 μl/well of Immunoprecipitation lysis buffer (Pierce IP Lysis Buffer^®^, Thermo Scientific), supplemented with Complete EDTA-Free Protease Inhibitor Cocktail tablet (04693159001, Roche Applied Science). When indicated, samples were treated with 100 μg/mL RNAseA for 20 minutes before co-immunoprecipitation. All lysates were cleared by centrifugation at 12.000 rpm for 30 min at 4°C. After protein quantification, lysates were incubated at a ratio of 1mg lysate/50μL of resin (mouse monoclonal anti-HA antibody (clone HA-7) conjugated to agarose beads, A2095, Sigma-Aldrich), rotating overnight at 4°C. Beads were then washed 4 times with IP Lysis Buffer and 2 times with PBS. For IP:IB experiments, co-immunoprecipitated proteins were recovered with one volume of 4X Laemmli sample buffer (Bio-Rad) containing 2-ME and for Mass-Spectrometry, with 200 μl of 250μg/mL HA-peptide (I2149, Sigma Aldrich) per 50 μl of resin. Aliquots of eluates were resolved by 10% SDS-PAGE, transferred to PVDF membranes (EMD Millipore) and blocked for 1h at RT with 3% non-fat dry milk in PBS containing 0.1% Tween-20, Membranes were probed with the desired primary antibody (incubated overnight at 4°C), followed by incubation with HPR-conjugated antibody (1h at RT) and visualized using SuperSignal^TM^ West Pico PLUS Chemiluminescent Substrate (34580, Thermo Scientific). Alternatively, gels were stained with SilverQuest^TM^ Silver Staining Kit (LC6070, Thermo Scientific), following manufacturer’s instructions.

### Mass spectrometry

Proteins present in eluates were concentrated using Ultra-4, membrane PLGC Ultracel-PL (UFC801024, Amicon) and resuspended in TNE (50 mM Tris pH 8.0, 100 mM NaCl, 1 mM EDTA) buffer. Samples were adjusted to 0.1% RapiGest SF reagent (Waters Corp.) and boiled for 5 min, followed by addition of TCEP (Tris (2-carboxyethyl) phosphine) to 1 mM final concentration and incubation at 37°C for 30 min. Samples were carboxymethylated with 0.5 mg/ml of iodoacetamide for 30 min at 37°C, followed by neutralization with 2 mM TCEP, and digested with trypsin (trypsin:protein ratio - 1:50) overnight at 37°C. RapiGest was degraded and removed by treatment with 250 mM HCl at 37°C for 1 h, followed by centrifugation at 14,000 rpm for 30 min at 4°C. The soluble fraction was applied to a C18 desalting column (Thermo Scientific, PI-87782). Desalted peptides were eluted from the C18 column into the mass spectrometer using a linear gradient (5–80%) of ACN (Acetonitrile) at a flow rate of 250 μl/min for 1h. The buffers used to create the ACN gradient were: Buffer A (98% H_2_O, 2% ACN, 0.1% formic acid, and 0.005% TFA) and Buffer B (100% ACN, 0.1% formic acid, and 0.005% TFA). Analysis of desalted-peptides was performed by ultra high-pressure liquid chromatography (UPLC) coupled with tandem mass spectroscopy (LC-MS/MS) using nano-spray ionization. Nano-spray ionization was done using a TripleTof 5600 hybrid mass spectrometer (ABSCIEX) interfaced with nano-scale reversed-phase UPLC (Waters corporation nano ACQUITY) using a 20 cm-75 micron ID glass capillary packed with 2.5-μm C18 (130) CSH^TM^ beads (Waters corporation). MS/MS data were acquired in a data-dependent manner in which the MS1 data was acquired for 250 ms at m/z of 400 to 1250 Da and the MS/MS data was acquired from m/z of 50 to 2,000 Da. The Independent data acquisition (IDA) parameters were as follows; MS1-TOF acquisition time of 250 milliseconds, followed by 50 MS2 events of 48 milliseconds acquisition time for each event. The threshold to trigger MS2 event was set to 150 counts when the ion had the charge state +2, +3 and +4. The ion exclusion time was set to 4 seconds. Finally, the collected data were analyzed using Protein Pilot 4.5 (ABSCIEX) for peptide identifications. Identified proteins were considered specific when at least two or more unique tryptic peptides were detected with a degree of confidence of 99% [69], and were never present in HA-USP14 or 3rLCMV-HA-GFP negative controls. Spectral count normalization (NSC) was used to estimate the relative protein abundance as described in [70]. Hits identified in 4 independent experiments were ranked as depicted in Table 1.

### siRNA

A549 cells were transfected (in triplicate) with 100nM siRNA (siGENOME Smartpool, Dharmacon) directed to each of the 12 selected hits listed in Table 1, using Hi-Perfect reagent (Qiagen) and media was replenished 6 hours post-transfection, according to manufacturer’s protocol. As control, cells were transfected with scrambled siRNA Pool 1 (Scr1) or Pool 2 (Scr2) (Dharmacon). siRNA transfection efficiency was evaluated using siGLO RNAi control (Dharmacon) and flow cytometry (FITC channel).

### DDX3-knockout A549 cell lines

DDX3 ko-1 and DDX3 ko-2 cell lines were generated by CRISPR/Cas9-mediated genome engineering following the protocol and algorithm described by [67] A target sequence in the first (DDX3 ko-1) or fifth (DDX3 ko-2) of human DDX3 was chosen and appropriate oligonucleotides were cloned into the BbsI site of pSpCas9(BB)-2A-GFP plasmid. (Primers: DDX3X-Exon1-Fw: CACCGAGTGGAAAATGCGCTCGGGC, DDX3X-Exon1-Rv: AAACGCCCGAGC GCATTTTCCACTC and DDX3X-Exon5-Fw: CACCGCGGAGTGATTACGATGGCAT, DDX3X-Exon5-FRv: AAACATGCCATCGTAATCACTCCGC. The plasmid was transfected for 24 hours, and the GFP-positive population was sorted by single-cell flow cytometry on a 96 well culture plate using a BD FACS Aria II Cell-Sorter. As control, A549 were transfected with empty plasmid (WT-pCas9 cells). Cells were expanded, maintained for a minimum of ten passages before their use and tested for DDX3 expression by Immunoblot with the aforementioned anti-DDX3 Ab.

### Cell viability

A549 cell viability was evaluated after knockdown with different siRNAs and after stable ko of DDX3 gene, prior to viral infection with LCMV, using Ghost-dye (Tombo Biosciences) and analysed using a BD LSRII Cytometer and FlowJo software (Treestar, Inc., Ashland, OR, USA).

### Retroviral mediated DDX3 reconstitution

DDX3 gene was cloned into pMD145 vector and retrovirus was assembled in Phoenix-AMPHO (ATCC^®^ CRL-3213^TM^) retrovirus packaging cell line after transfection with TransIT-293 transfection reagent (Mirus Bio LLC). The empty vector was also used as a negative control. After 48 hours, supernatant was collected, filtered and used for A549 transduction using 20μg/mL DEAE/Dextran. Plates were centrifuged at 1200xg for 40 minutes at room temperature and transduced cells were selected with 1.5μg/mL Puromycin.

### Quantitative image-based analysis

Virus-infected cells were fixed in 10% buffered formalin for 72 h and blocked in 3% bovine serum albumin-PBS for 1 h. Cells were then stained with murine mAbs against JUNV or LASV glycoprotein (GD01, L-52-161-6 antibodies, respectively, 1:1,000 dilution in blocking solution), followed by Alexa Fluor 488-conjugated goat anti-mouse IgG (ThermoFisher) (1:1,000 dilution in blocking solution). All infected cells were also stained with Hoechst 33342 and HCS Cell Mask Red (ThermoFisher) for nuclei and cytoplasm detection, respectively. High-content quantitative imaging data were acquired and analyzed on an Opera confocal reader (model 3842 and 5025; quadruple excitation high sensitivity; Perkin-Elmer), at two exposures using a ×10 air objective lens as described previously [71]. Analysis of the images was accomplished within the Opera environment using standard Acapella scripts. Nuclei and cytoplasm staining were used to determine total cell number and cell borders, respectively. Mock-infected cells were used to establish a threshold for virus-specific staining. Quantification of virus positive cells was subsequently performed based on mean fluorescent intensities in the virus-specific staining channel. Infection rates were then determined by dividing the number of virus positive cells by the total number of cells measured.

### qPCR

For BSL-2 analyses total RNA was extracted using RNeasy kits (Qiagen), and reverse transcribed into cDNA using MMLV RT (Invitrogen). cDNA quantification was performed using SYBR Green PCR kits (Applied Biosystems) and a Real-Time PCR Detection System (ABI). For BSL-4 infections, viral RNA yields from the media were determined by qRT-PCR as previously described [71]. Briefly, RNA was extracted with Trizol (Thermo Fischer Scientific) and the Ambion Blood RNA Isolation Kit (Thermo Fischer Scientific). The assay was performed with RNA Ultra Sense one-step kit (Thermo Fisher Scientific) and TaqMan Probe (ABI, Thermo Fischer Scientific) following the manufacturer’s instructions. The primers used were: LCMV-GP-Fw: CATTCACCTGGACTTTGTCAGACTC, LCMV-GP-Rv: GCAACTGCTGTGTTC CCGAAA, LCMV-NP-Fw: GCATTGTCTGGCTGTAGCTTA, LCMV-NP-Rv: CAAT GACGTT GT ACAAGCGC; JUNV-NP-Fw: CGCCAACTCCATCAGTTCATC, JUNV-NP-Rv: CCATGAGGAGTGTTCAACGAAA; probe JUNV NP Prb: 5-6FAM-TCCCCAGATCTCCCACCTTGAAAACTG-TAMRA; LASV-GPC-Fw: GCAGTGCTG AAAGGTCTGTACAA, LASV-GPC-Rv: AGGAGGAAAGTGACCAAACCAA, probe LASV-GPC: 5-6F AM-TTT GCAACGT GT GGCCT-T AMRA; SeV-NP-Fw: TGCCCTGGAAGATGAGTTAG, SeV-NP-Rv: GCCTGTTGGTTTGTGGTAAG; huIFNb Fw: AAACTCATGAGCAGTCTGCA, huIFNb-Rv: AGGAGATCTT CAGTTTCGGAGG huGAPDH-Fw: T GAT GACAT CAAGAAGGT GGT GAAG and huGAPDH-Rv: TCCTTGGAGGCCATGTGGGCCAT. Serial 10-fold dilutions of the assayed (10^2^ to 10^7^ copies) virus were used as standards. Relative expression levels were determined by using the comparative cycle threshold method.

### Bioactive IFN-I

Human IFN-I bioactivity in tissue culture supernatants was measured with reference to a recombinant human IFN-β standard (InvivoGen) using HEK-Blue^TM^ IFN-α/β cell line (InvivoGen) and QUANTI-Blue^TM^ detection reagent, following manufacture’s instructions.

### Minireplicon and Translation assay

LCMV minireplicon system was assayed as described elsewhere [72]. Briefly, WT and DDX3 ko A549 cells were transiently cotransfected, using Lipofectamine 2000, with 0.6 μg of pCAGGS L, 0.15 μg of pCAGGS NP and 0.5 μg of the dual-reporter (green fluorescent protein (GFP) and *Gaussia* luciferase (Gluc)) minigenome (MG) plasmid. These constructs were driven by the human polymerase-I promoter [73]. To normalize transfection efficiencies, 0.1 μg of a mammalian expression vector encoding *Cypridina noctiluca* luciferase (Cluc) under the control of the constitutively active simian virus 40 (SV40) promoter (pSV40-Cluc; New England BioLabs), were included in the transfection mix. GFP expression was determined by fluorescence microscopy using a Leica fluorescence microscope. Microscope images were pseudocolored using Adobe Photoshop CS4 (v11.0) software and by luminometry (Gluc) using a Lumicount luminometer (Packard). Cells were also subjected to flow cytometry analysis at 72 h post-transfection, and percentages of GFP-positive (GFP+) cells and mean fluorescence intensities (MFI) of the FL1-gated cell population were determined using FlowJo software (Tree Star). Luciferase gene activities were determined using Biolux *Gaussia* and *Cypridina* Luciferase Assay kits (New England BioLabs) using a Lumicount luminometer (Packard). Reporter gene activation (Gluc) is indicated as fold induction over cells transfected with a negative pCAGGS empty plasmid control instead of the viral NP. The translation assay was performed as indicated in [34]. Capped synthetic RNAs were obtained by *in vitro* transcription from T7 promoter-controlled constructs. The virus-like mRNA (5’wt/3’wt_2), mimicking the TCRV NP mRNA comprises a 5-nt nonviral sequence preceding the viral 5’UTR, which is fused to the reporter firefly Luciferase (FLUC) open reading frame followed by the viral 3’ UTR. The cell-like 5’βGlo/3’poly(A) transcript bears the 5’UTR from human β-globin, and a 53-nt 3’ poly(A) tail flanking the FLUC coding sequence. Briefly, HEK-293T cells were grown in 24 well plates, transfected with 50 pmol of siRNAs against DDX3 or scrambled siRNA pool 1 (Scr), and 42 hours later transfected again with 200 ng per well of the indicated capped synthetic RNA. As internal control, 75 ng/well of a Renilla Luciferase (RLUC)-expressing non-capped mRNA was added to the transfection mix. Following 6h incubation, lysis of transfected cells and quantification of FLUC and RLUC activities on a Biotek FLx800 luminometer were performed using Dual-luciferase reporter assay system (Promega), according to the manufacturer’s instructions. FLUC activity was normalized against the corresponding value of RLUC activity in each experimental condition. For each transcript, mean FLUC values (+/− standard deviation; SD) determined in depleted cells, are shown as a percentage of those in control cells, taken as 100%.

### Statistics

Statistical differences were determined by Student’s t test or by one-way or twoway analysis of variance (ANOVA) followed by Bonferroni post-hoc analysis using the GraphPad Prism 5 software (La Jolla, CA). For the Translation assay, statistical analyses were performed using the SPSS 17.0 statistical software package (SPSS, Inc., Chicago, IL, USA).

### Data availability

Mass spectrometry results were deposited in http://www.peptideatlas.org/PASS/PASS01114.

### Ethics statement

This study was carried out in strict accordance with the recommendations in the Guide for the Care and Use of Laboratory Animals of the National Institutes of Health under a protocol (S07315) approved by the Institutional Animal Care and Use Committee at the University of California, San Diego (Animal Welfare Assurance Number: D16-00020). All efforts were made to minimize suffering of animals employed in this study.

## ACKNOWLEDGEMENTS

We are deeply grateful to Dr. Ellen Wehrens for very insightful feedback on our manuscript and to Dr. Eric Bennett (Division of Biological Sciences, UCSD) for thoughtful advice in designing Mass-Spectrometry experiments and providing HA-USP14 plasmid (PMID: 19615732). Plasmids encoding LASV, JUNV, MACV and TCRV NPs were generously provided by Dr. Luis Martinez-Sobrido (Rochester Medical Center, NY. USA). Plasmid encoding DDX3 was a gift from Dr. Ricardo Soto-Rifo (Facultad de Medicina, Universidad de Chile) and plasmid pMD145 and Phoenix-AMPHO cells, from Dr. Matthew Daugherty (Division of Biological Sciences, UCSD). pSpCas9(BB)-2A-GFP plasmid was obtained from the laboratory of Dr. Feng Zhang (McGovern Institute for Brain Research at MIT) via Addgene (I.D. 48138). Sendai Virus was kindly provided by Dr. Carolina López (Penn Institute for Immunology, University of Pennsylvania).

